# A field-capable rapid plant DNA extraction protocol using microneedle patches for botanical survey and monitoring

**DOI:** 10.1101/2023.01.09.523204

**Authors:** Jonathan Selz, Nicolas R. Adam, Céline E. M. Magrini, Fulvia Malvido Montandon, Sven Buerki, Sebastian J. Maerkl

## Abstract

**Premise:** A novel protocol for rapid plant DNA extractions using microneedles is proposed, which supports botanic surveys, taxonomy and systematics. This protocol can be conducted in the field with limited laboratory skills and equipment. The protocol is validated by conducting sequencing and comparing results with Qiagen spin-column DNA extractions using BLAST analyses.

**Methods and Results:** Two sets of DNA extractions were conducted on 13 species spanning various leaf anatomies and phylogenetic lineages: i) fresh leaves were punched with custom polymeric microneedle patches to recover genomic DNA ii) Qiagen DNA extractions. Three plastid (*matK*, *rbcL*, *trnH-psbA*) and one nuclear ribosomal (ITS) DNA regions were amplified, and Sanger or Nanopore sequenced. The proposed method reduced the extraction to 1 min and yielded the same DNA sequences as Qiagen.

**Conclusions:** Our drastically faster and simpler method is compatible with Nanopore sequencing and suitable for multiple applications including high-throughput DNA-based species identification and monitoring.

## INTRODUCTION

The rising human population puts ever-growing pressure on natural ecosystems through deforestation, natural resource exploitation, and climate change. Earth’s biodiversity is disappearing at an alarming rate, placing ecosystems and their services in jeopardy (Antonelli et al., 2020). These services are crucial to support life as we know it, including factors such as food production, medicinal resources, materials and more broadly the regulation of climate, the formation of fertile soil, and maintaining water and air quality. The protection of nature is crucial to retain its economical and societal benefits (Almond et al., 2020). Biodiversity identification and monitoring are the cornerstones for effective preservation strategies. However, the time required by classical identification approaches no longer fit the dynamics of ecosystem degradations. Species are disappearing at a higher rate than they are being identified. To tip the scales of this race against time, initiatives have been put in place to improve the speed of biodiversity assessment (Conservation International, 2022) and to create a database of standardized DNA barcodes for faster species identification (International Barcode of Life, 2022).

These DNA barcodes are produced by extracting DNA using well-established protocols such as CTAB or spin-column extractions and amplifying the targeted DNA regions with Polymerase Chain Reaction (PCR). Sequencing techniques, such as Sanger, are then used to generate the barcoding data from these amplicons. This lengthy process relies heavily on laboratory equipment and specialized skills. Next-generation sequencing (NGS) brought promising prospects for DNA-based species identification (Buerki and Baker, 2016) and improvements in sequencing technology was shown to enable on-site species identification in remote environments with a mobile laboratory (Pomerantz et al., 2018). However, these genomic methods suffer from lengthy and cumbersome DNA extraction and sample preparation processes, particularly for plants, which impede efficient field protocols. Polymeric microneedle (MN) patches have been initially developed for drug delivery (Larrañeta et al., 2016) and point-of-care diagnostic (He et al., 2020). Recent studies have shown MN patches to be a promising method for rapid DNA extraction with applications for the detection of allergens in food matrices (Li et al., 2021) and for plant pathogen detection (Paul et al., 2019). These DNA extractions using MN patches require virtually no equipment or laboratory skills and take less than a minute to execute. In addition, this non-destructive method does not require large amounts of plant tissue or lengthy processing steps, such as grinding, and is therefore a promising option for fragile or scarce sources such as herbarium samples. However, quantifications of MN extractions were shown to yield lower DNA concentrations and purity than methods such as CTAB (Paul et al., 2019). Also, it is still unclear whether MN DNA extractions can be used to extract host plant DNA and especially nuclear DNA. Achieving this would allow using this DNA extraction technique to support rapid botanic surveys, taxonomy, and systematics by reducing the time, cost, and skill barriers associated with genomic methods. Here, we propose a new protocol to test this approach and compare the results with spin-column DNA extraction.

In this study, MN patches were shown to be able to extract nuclear DNA from plant samples for species identification using multi-locus plastid and nuclear DNA barcoding. The proposed protocol was developed in the frame of GenoRobotics, an EPFL interdisciplinary project involving students, engineers, and researchers, and tested on plants of the EPFL campus. The objectives of this study are (1) to develop a custom fabrication process for MN patches using standard laboratory equipment, (2) to develop a rapid and field-deployable DNA extraction protocol using MN patches, (3) to sequence not only plastid but also nuclear DNA regions from MN DNA extractions. To achieve these objectives, a cost-efficient alternative to commercially available MN molds such as those offered by Blueacre Technology (Blueacre Technology Ltd., Dundalk, Ireland) was investigated to reduce the cost of producing the master molds. Further, the quality of MN DNA extraction must enable downstream PCR amplification and Sanger or next-generation sequencing using Nanopore technology. To ensure this quality and since MN DNA extraction’s purity was shown to be limited, a DNA purification step was added. We then employed a multi-locus approach with the widely used plastid markers *matK*, *rbcL*, *trnH*-*psbA* and the nuclear ribosomal ITS region (including ITS1, 5.8S and ITS2) (Hollingsworth, 2011). We also applied the same protocols to DNA extracted with a standard spin-column based method. Sequencing results from both DNA extraction methods were validated against each other by conducting BLAST similarity analyses (Morgulis et al., 2008) to ensure that the same sequences were produced. These sequencing results were then matched against the NCBI database to confirm that the correct DNA regions were sequenced, and that the same species were obtained. Finally, the applicability of MN extraction to Nanopore sequencing was evaluated on 2 species.

## METHODS AND RESULTS

### Materials

Custom MN patches were produced in a three-step process (Figure 1A-B): printing master molds, using the master molds to create PDMS molds, and casting MN patches using PDMS molds (Wang et al., 2017). MN patch geometry is a 1cm^2^ array of 121 conical needles with a height of 1600 μm. The material of the MN patches is Poly(vinyl alcohol) hydrogel. Master molds were 3D printed by the Atelier de Fabrication Additive at EPFL in HTM 140 resin with the Perfactory^®^ 4 Mini XL 3D printer (EnvisionTec Inc, Dearborn, USA) using SLA/DLP technology. This method has a resolution in the order of tens of microns. Theses master molds were then used to cast PDMS molds with a 1:10 mix of hardener to elastomer of Sylgard 184 (Suter Kunststoffe AG, Fraubrunnen, CH). A solution of H_2_O (UltraPure^™^ DNase/RNase-Free Distilled Water; Thermo Fisher Scientific Inc., Waltham, USA) and Poly(vinyl alcohol) (Merck & Cie, Schaffhausen, CH) with a 7:1 weight ratio was prepared. After cleaning the PDMS molds in a heated bath of bi-distilled water, letting them dry, and placing them in 12-well plates, 750 μL of PVA solution was added to each well. The plates were then centrifuged at 2900 × g (4000 rpm) for 20 min at 40°C. Another 200-750 μL of the PVA solution was added to fill the molds, taking care not to overfill them. After a drying time of 36 hours in a fume hood, the MN patches were unmolded and ready for use.

**FIGURE 1:**
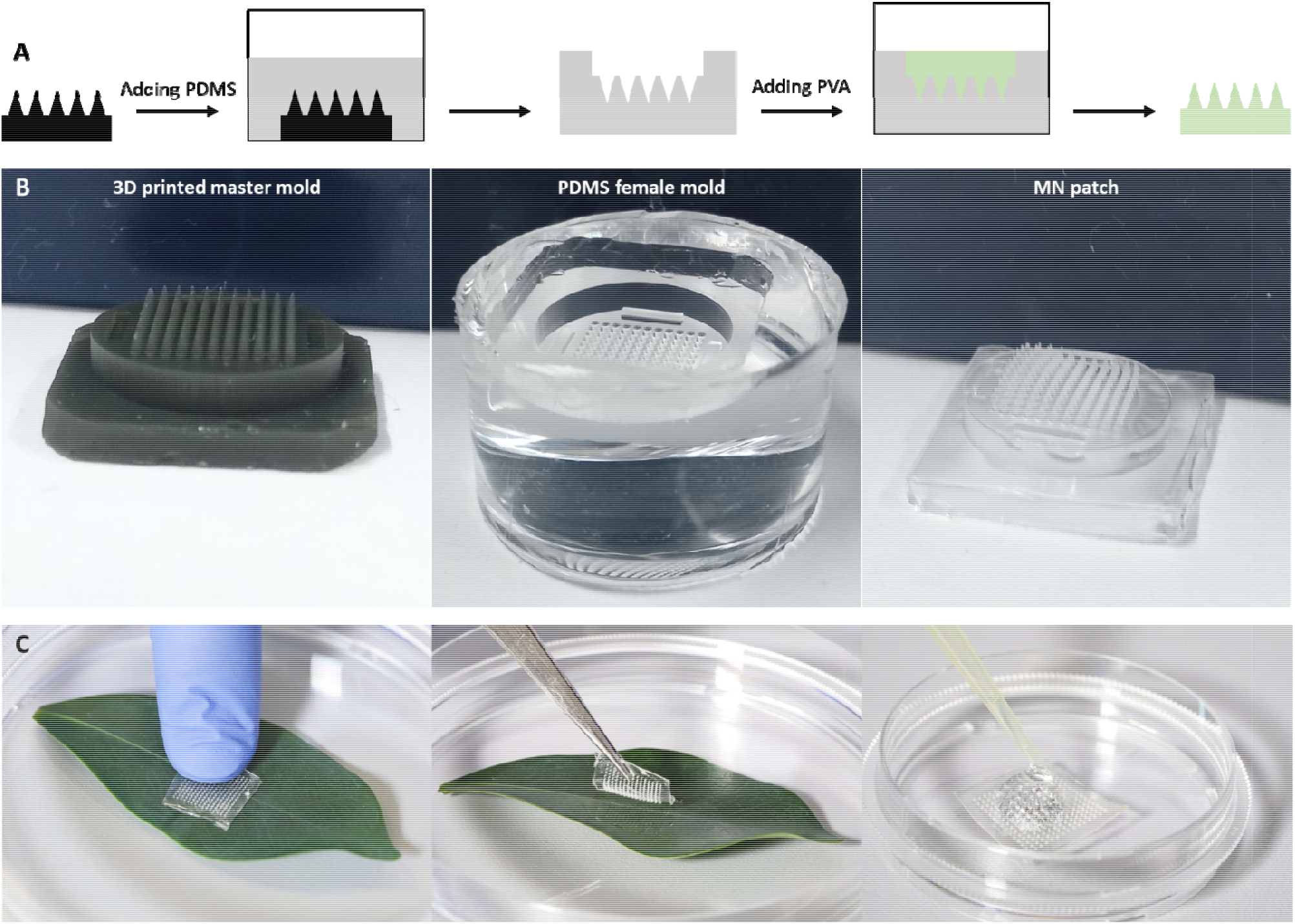
MN patch fabrication (**A-B**)and usage (**C**). **A**, Schematic of the fabrication steps from the 3D printed master mold to the final polymeric MN patch. **B**,The three main steps in the MN fabrication process. **C**,Demonstration of the DNA extraction process consisting of punching a leaf sample with the MN patch and eluting the DNA.

Initially, *Solanum lycopersicum* L. was used as a model to design and test the DNA extraction and amplification protocols as well as to test the MN fabrication process. In a second phase, the protocol was tested on additional plants to validate its applicability to phylogenetic lineages. Eight species were sampled around the EPFL campus (46°31’11.050”N, 6°33’57.966”E) and five species were purchased and grown in the laboratory. All selected plants are angiosperms and Eudicots except for one specimen of the Poales order, which belongs to Monocots. The selection of samples for this study was based on targeting a wide diversity of plants in order to test the MN patches on different leaf morphologies. The notable variations are leaf thickness, succulence and fibrousness; presence and thickness of cuticle; presence of latex, oils and mucilage. The availability of a reference sequence was also a critical selection criterion for the sampled species. The plant samples and their taxonomy are summarized in Appendix 1.

### MN DNA extractions

Genomic DNA was extracted from fresh leaf material (Appendix 1). 2-3 young leaves were selected and stacked to ensure a sufficient thickness compared to the height of the microneedles. A MN patch was then used to puncture the leaves by placing it in an area with as few veins as possible and pressing the patch forcefully against the leaf between the thumb and index for 10-15 sec. DNA was then recovered by eluting the samples from the MN patches with 50 μL of H_2_O (UltraPure^™^ DNase/RNase-Free Distilled Water; Thermo Fisher Scientific Inc., Waltham, USA). This critical step requires wetting of the full surface of the MN patch, by pipetting the buffer up and down until bubbles form. It is important not to let the buffer sit for too long to avoid dissolution of the patch itself. The full extraction process (Figure 1C) requires about 1 minute to be completed. All MN extractions were then purified with DNA purification columns according to the manufacturer’s protocol (QIAquick; Qiagen AG Hombrechtikon, CH). We used 50 μL of extracted DNA as input and the final elution was performed with 30 μL of H_2_O.

### Qiagen DNA extractions

DNA extraction with spin columns (DNeasy Plant Mini Kit; Qiagen AG, Hombrechtikon, CH) was selected as a reference method for its relative simplicity, speed, reliability, and widespread usage. Leaves were sampled and dried in silica gel for 36 hours. Leaf tissue was then finely ground with a mortar and pestle and 20 mg of dried leaf powder was used as an input for the DNA extraction protocol, which was carried out according to the manufacturer’s protocol. Final elution was performed in 2 steps of 50 μL each and pooled together.

### DNA amplification and sequencing on both DNA extraction methods

Four commonly used DNA barcode regions in plants (Hollingsworth, 2011) were selected: the coding regions of the *matK* and *rbcL* plastid genes (*matK* and *rbcL*), the *trnH-psbA* plastid intergenic spacer, and the ITS region of nuclear ribosomal DNA. The primers used were taken from the following sources: matK472F and matK1248R (Yu et al., 2011), rbcLa-F (Kress and Erickson, 2007) and rbcL724R (Fay et al., 1997), trnH (Tate, 2002) and psbA (Sang et al., 1997), ITS-p5 and ITS-p4 (Cheng et al., 2016). Amplifications of these four target DNA regions were carried out in 50 μL reactions, containing 25 μL of 3 mM MgCl_2_ PCR master mix (Taq 2X Master Mix; New England Biolabs Inc., Ipswich, USA), 1 μL of each primer at 10 μM stock concentration as well as 16 μL H_2_O, and 7 μL of DNA template for the MN extractions or 21 μL H_2_O and 2 μL of DNA template for the Qiagen extractions. The following PCR profile was used: initial denaturation at 95°C for 2 min, followed by 45 cycles of denaturation at 95°C for 30 sec, annealing at 54°C for 30 sec, extension at 68°C for 1 min, followed by a final extension at 68°C for 10 min. PCR products were then sent to a sequencing service (Microsynth AG, Balgach, CH) for purification and Sanger sequencing of both forward and reverse primers.

In addition to Sanger sequencing, the applicability of Nanopore sequencing to DNA extracted from MN patches was tested. PCR products for the four targeted DNA regions of two species, *Solanum lycopersicum* L. and *Ficus benjamina* L., were sequenced using a MinION Mk1B (Oxford Nanopore Technologies, Oxford, United Kingdom). Oxford Nanopore Technologies (ONT) offers two types of Flow Cells for the MinION. The Spot On Flow Cell Mk 1 R9 Version 1 (FLO-MIN106D) has higher sequencing capabilities compared to the Flongle Flow Cell (FLO-FLG001) and can therefore generate more sequencing reads but is ten times more expensive. *Solanum lycopersicum* L. samples were sequenced using MinION and Spot On Flow Cell. *Ficus benjamina* L. was sequenced using a Flongle Flow Cell with a Flongle adapter for MinION (ADP-FLG001) for further cost reduction of the process. Nanopore library preparation of all samples was carried out with the Rapid Barcoding Kit (SQK-RBK004). The base calling is performed with the MinKNOW software (Oxford Nanopore Technologies, Oxford, United Kingdom), which generates Fast5 and Fastq files containing all the sequencing reads.

### Analyses

To assess the validity of the MN DNA extractions, we compared DNA sequencing results obtained from MN extracted samples to sequencing results obtained from standards extracted with the Qiagen method. The sequencing output composed of forward and reverse sequences were aligned using MAFFT v7 (Katoh and Standley, 2013) on the ab1 files with a minimum quality level of 20 to obtain the consensus sequence of the amplified DNA region. When the overlap between the forward and reverse sequences was too small, consensus sequences were obtained by manually aligning the fragments on the reference sequence from the NCBI database (Table 1). To ensure that both methods produced the same sequencing data, a first validation was performed by aligning each MN sequence on its Qiagen counterpart using BLAST 2 seq (Zhang et al., 2000). These sequences were then searched on GenBank’s Nucleotide database using MegaBLAST (Morgulis et al., 2008) to evaluate if the quality of the sequences enabled species identification (Table 1; Figure 2). The comparison metrics were the identity percentage (IP), which indicates the similarity between the query and the reference sequence, and the query coverage (QC), which is the percentage of the query length aligned on the reference. The analysis of the Nanopore sequences was performed by aligning the Fastq reads on a reference sequence. DNA sequence alignments were obtained using Fastq Custom Alignment from Epi2me software (Oxford Nanopore Technologies, Oxford, United Kingdom) with a minimal Q-score of 7.

**TABLE 1.**
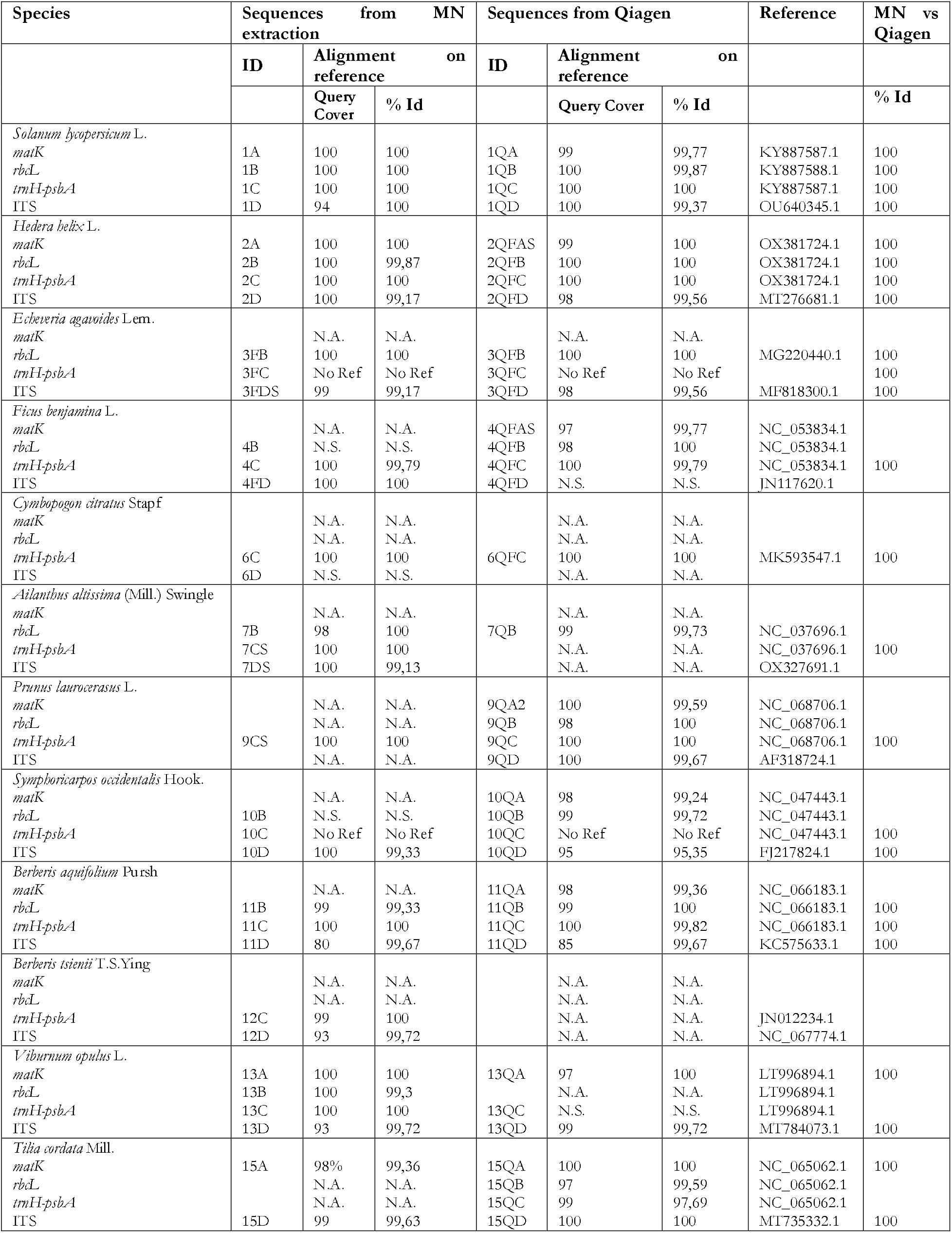
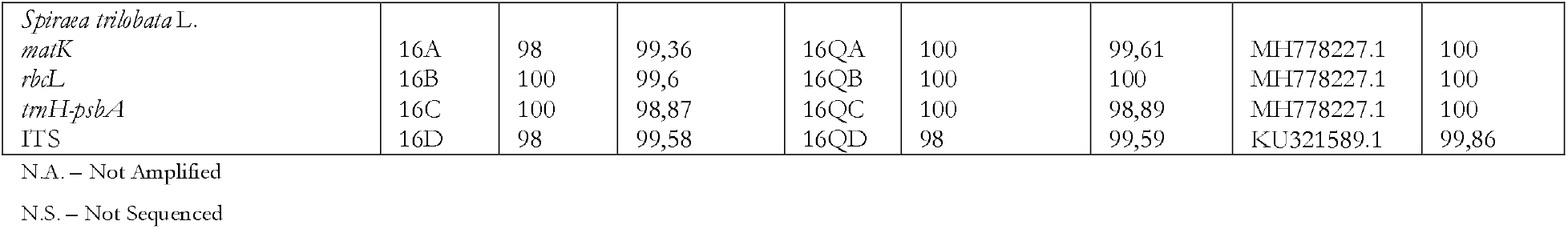
Summary of the Sanger sequences’ alignment against the reference sequences of GenBank’s Nucleotide database on DNA extracted with MN and Qiagen.

**FIGURE 2:**
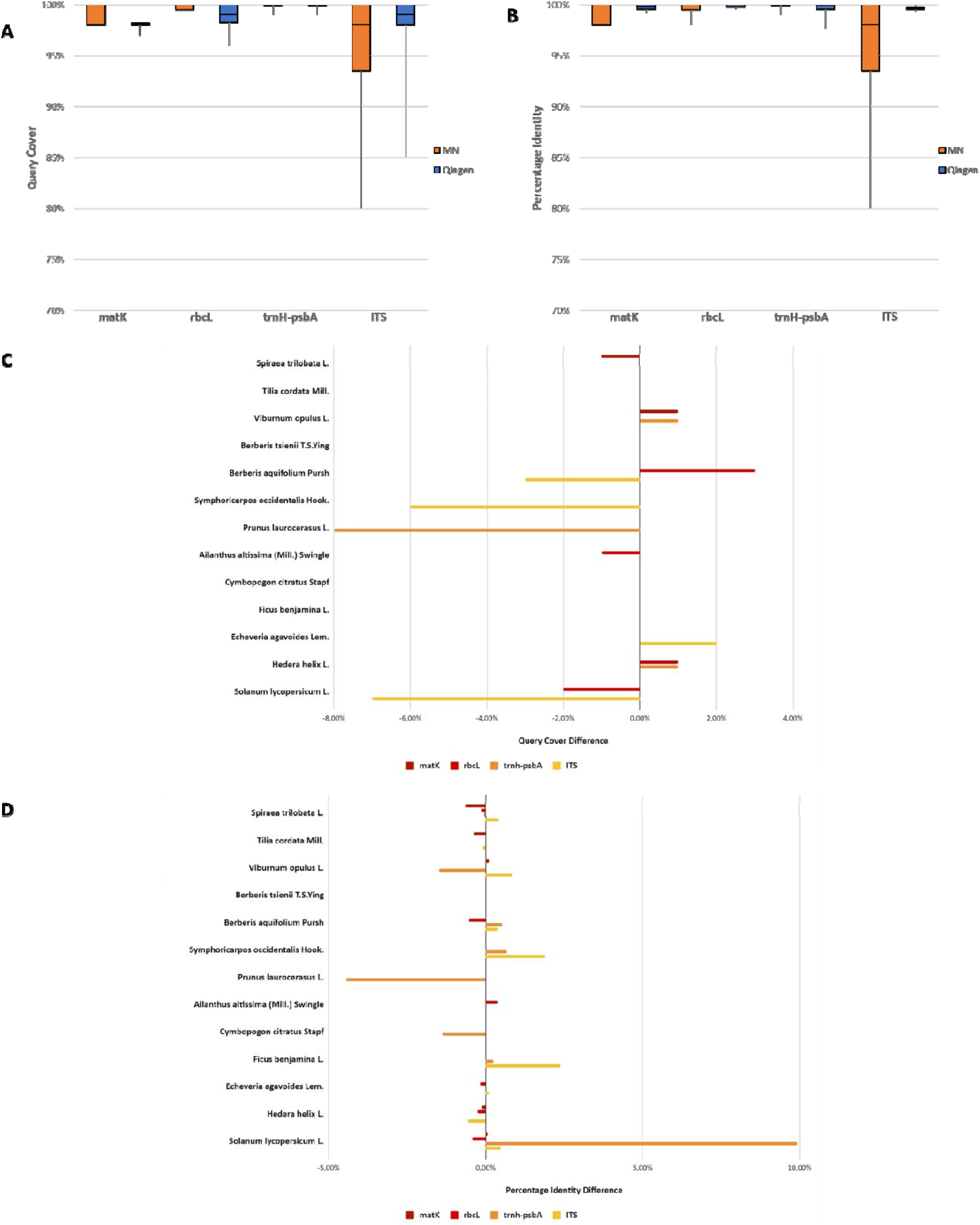
MN and Qiagen sequence alignments results of the 4 DNA regions of the 13 plant samples, the boxplots span the first quartile to third quartile and the whiskers indicate the lowest values obtained. **A**, Distribution of the Query Cover for the alignments of the different DNA regions against the NCBI database. **B**, Distribution of the Percentage Identity for the alignments of the different DNA regions against the NCBI database. **C**, Query Cover difference obtained by the subtraction of Qiagen to MN results. **D**, Percentage Identity difference obtained by the subtraction of Qiagen to MN results

### Results

Out of the four targeted DNA regions sequenced for 13 plant species (i.e., a total of 52 DNA sequences), 34 successful sequences were obtained from MN DNA extractions versus 39 from Qiagen extractions. Unsuccessful amplifications of the targeted DNA regions were the main cause of failure. For *matK*, 38.5% of MN samples against 69.2% of Qiagen samples yielded successful sequences. For *rbcL*, sequences were obtained for 53.8% of MN samples against 76.9% for Qiagen. For *trnH*-*psbA* and ITS, the process was successful for 84.6% of the MN samples against 76.9% for the Qiagen samples. Considering the stochasticity on the total number of sequenced samples, both methods performed similarly. The success rate for *matK* is significantly lower than for the other DNA regions. The amplification of the *matK* region is notably more susceptible to the sample’s purity and could be improved for instance through the addition of dimethyl sulfoxide (Buerki et al., 2009).

The BLAST analyses performed on the sequencing data from MN and Qiagen extractions showed a 100% similarity between MN and Qiagen sequences (Table 1). The MN DNA extraction method can therefore be used to generate the same sequencing data as standard Qiagen extractions. A notable exception is *trnH-psbA* for *Prunus laurocerasus* L., for which the MN sample yielded very short sequences. The BLAST comparison against GenBank’s database (Table 1; Figure 2) demonstrated that the generated sequences correspond to the targeted DNA regions of the sampled species. The top hits in the database were the same for both MN and Qiagen methods, validating the applicability of MN DNA extractions for species identification. The consistency of the IP for the MN sequences also indicates the high repeatability of this method. It is also interesting to note the high IP and QC of the MN method for the ITS region, showing that nuclear DNA can be obtained reliably using this extraction method. For *Ficus benjamina* L., the plastid DNA regions and the ITS region seems to yield two different *Ficus* species, hence the taxonomy could be validated only at the genus level. The Nanopore sequencing results (Table 2) showed an excellent similarity between Qiagen and MN methods for both *Solanum lycopersicum* L. and *Ficus benjamina* L. The minimal average identity is 95.6% for MN sequences and 96.3% for Qiagen sequences when the minimal average accuracy is a bit higher for MN sequences than Qiagen sequences, respectively 88.6% and 79%. These results confirm the compatibility of MN DNA extractions with NGS and more specifically Nanopore sequencing.

**TABLE 2.** Summary of the Nanopore sequences’ alignment against the reference sequences of GenBank’s Nucleotide database on DNA extracted with MN and Qiagen from *Solanum lycopersicum* L. and *Ficus benjamina* L. samples.

| Species | MN Samples |  |  | Qiagen Samples |  |  | Reference |
| --- | --- | --- | --- | --- | --- | --- | --- |
|  | ID | FASTQ custom alignment |  | ID | FASTQ custom alignment |  |  |
|  |  | Average accuracy | Average identity |  | Average accuracy | Average identity |  |
| *Solanum lycopersicum* L. |  |  |  |  |  |  |  |
| *matK* | 1A | 91,2 | 97,1 | 1QA | 93,1 | 97,1 | KY887587.1 |
| *rbcL* | 1B | 91,4 | 96,8 | 1QB | 93,7 | 96,8 | KY887588.1 |
| *trnH-psbA* | 1C | 91,2 | 97,5 | 1QC | 93,2 | 97,5 | KY887587.1 |
| ITS | 1D | 88,6 | 95,6 | 1QD | 83,5 | 99,6 | OU640345.1 |
| *Ficus benjamina* L. |  |  |  |  |  |  |  |
| *matK* |  | N.A. | N.A. | 4QA | 93 | 97,6 | NC_053834.1 |
| *rbcL* | 4B | 92,2 | 96,9 | 4QB | 79 | 97,2 | NC_053834.1 |
| *trnH-psbA* | 4C | 92,4 | 97,6 | 4QC | 99 | 97,5 | NC_053834.1 |
| ITS | 4D | 90 | 97,5 | 4QD | 89,3 | 96,3 | JN117620.1 |

N.A. – Not Amplified

The previous results show that both extraction methods produce similar sequencing data, however spin column and microneedle-based DNA differ heavily in the way they are carried out. While the Qiagen spin column extraction requires a thermal heating/cooling block, a centrifuge, several reagents and consumables, the MN extraction is performed with only a single MN patch, pipet and tip, 1.5 mL tube and H_2_O. MN extraction is a straightforward process that requires no laboratory know-how. The process is therefore cost and time effective. MN extraction can be performed in 1 min plus ~ 10 min for purification, while a Qiagen extraction requires at least 30 min for sample preparation and 90 min for the extraction protocol, hence more than 10 times longer. In terms of cost, the Qiagen kit costs about 4.50 USD per sample. The fabrication of one MN patch amounts to less than 0.30 USD and the purification costs 1.30 USD per sample, hence 2.75 times cheaper. MN extraction is drastically cheaper than commercially available DNA extraction kits especially with further improvements on the purification step. Overall, this new protocol has the potential to minimize the current biodiversity crisis by lowering the cost, time and skill barriers as well as the need for laboratory infrastructure when conducting DNA-based species identification in a field setting.

## CONCLUSIONS

We introduced a novel protocol to produce low-cost MN patches in-house with standard laboratory equipment using custom 3D printed master molds. We tested the DNA extractions using these MN patches on plants on campus and cultivated in the laboratory and compared them to Qiagen extractions. This study shows MN DNA extraction of genomic DNA from different plant species. With this method combined to a purification step, we managed to amplify and sequence plastid and nuclear DNA regions with Sanger and Nanopore technologies. The sequencing results showed the same outputs for both MN and Qiagen extractions as well as the usability of MN sequences for species identification through comparison with a reference database. This supports the application of the proposed technique for plant biodiversity survey, monitoring, taxonomy, and systematics.

The next step for a true field capable DNA analysis tool would be to overcome the need for thermal cycling during the amplification step by replacing PCR by an isothermal amplification method such as recombinase polymerase amplification and coupling it with Nanopore sequencing to analyze the amplicons. Additionally, the need for the DNA purification step needs to be further investigated. Avoiding this step would reduce the DNA extraction time even more and drastically improve field capability. Preliminary results seem to indicate a seasonal influence on the amplification output. Successful amplifications of unpurified MN DNA extractions were obtained in Spring when, in late Summer and Autumn, all amplifications without the purification step failed. Possible solutions could involve tuning the PCR parameters, or adapting the MN patch’s material by using a blend with a polycationic polymer (Kiang et al., 2004) or with a slightly cytotoxic one such as branched Polyethyleneimine (Zhang et al., 2011). Further investigations are however necessary to validate the impact of plant metabolism on MN DNA extractions and on the necessity of a purification step.

This protocol provides a simple and rapid method for sample collection in remote locations and large-scale studies for botanical survey and monitoring. MN DNA extraction provides plant DNA, which can be amplified and sequenced using Nanopore technology, and is 10 times faster and 2.75 times cheaper than conventional Qiagen extractions. Compared to commonly used extraction methods, its simplicity, its speed and its lack of lab equipment make it suitable for field DNA analysis and could serve as basis for high-throughput DNA-based species identifications and monitoring. Its ease of use combined with its low cost make it a first step towards citizen science, which would enable a high potential for data collection, and at the same time promote biodiversity awareness in local communities.

## ACKNOWLEDGMENTS

This work was supported by the EPFL School of Life Sciences, the School of Engineering, the Institute of Bioengineering, and the MAKE Fund. The authors would like to thank the Atelier de Fabrication Additive for the manufacturing of the Master Molds, the Jardin Botanique de Genève and particularly Beat Baumler for their field expertise, the EPFL DLL for the infrastructure and equipment, and finally Laura Kvedarauskaite for her initial work on the protocols.

## AUTHOR CONTRIBUTIONS

CEMM, FMM, JS and NRA conceived the protocols, performed the experiments and analyzed the data; CEMM, FMM designed the experiments; JS, NRA reviewed the experimental designs and wrote the manuscript; SJM and SB provided guidance and critical reviews. All the authors approved the final version of the manuscript

## DATA AVAILABILITY STATEMENT

Currently in submission (NCBI + Herbarium)

## Appendix 1. Plant samples and their taxonomy

N.A. – Not Amplified **Appendix 1.** Plant samples and their taxonomy
| ID | Species | Family | Order | Sampling location | Geographic coordinates | NCBI Accession |
| --- | --- | --- | --- | --- | --- | --- |
| 2 | *Hedera helix* L. | Araliaceae | Apiales | EPFL | Cultivated |  |
| 10 | *Symphoricarpos occidentalis* Hook. | Caprifoliaceae | Dipsacales | EPFL | 46°31'09.730"N<br>6°33'51.321"E |  |
| 13 | *Viburnum opulus* L. | Viburnaceae | Dipsacales | EPFL | 46°31'09.810"N<br>6°33'55.819"E |  |
| 15 | *Tilia cordata* Mill. | Malvaceae | Malvales | EPFL | 46°31'08.250"N<br>6°34'00.165"E |  |
| 6 | *Cymbopogon citratus* Stapf | Poaceae | Poales | EPFL | Cultivated |  |
| 11 | *Berberis aquifolium* Pursh | Berberidaceae | Ranunculales | EPFL | 46°31'09.339"N<br>6°33'51.022"E |  |
| 12 | *Berberis tsienii* T.S.Ying | Berberidaceae | Ranunculales | EPFL | 46°31'09.341"N<br>6°33'50.966"E |  |
| 4 | *Ficus benjamina* L. | Moraceae | Rosales | EPFL | Cultivated |  |
| 9 | *Prunus laurocerasus* L. | Rosaceae | Rosales | EPFL | 46°31'12.793"N<br>6°33'55.076"E |  |
| 16 | *Spiraea trilobata* L. | Rosaceae | Rosales | EPFL | 46°31'11.507"N<br>6°33'58.433"E |  |
| 7 | *Ailanthus altissima* (Mill.) Swingle | Simaroubaceae | Sapindales | EPFL | 46°31'11.290"N<br>6°33'58.347"E |  |
| 3 | *Echeveria agavoides* Lem. | Crassulaceae | Saxifragales | EPFL | Cultivated |  |
| 1 | *Solanum lycopersicum* L. | Solanaceae | Solanales | EPFL | Cultivated |  |

## Appendix 2. Required materials and protocol to perform the fabrication, DNA extraction and amplification steps

The protocol can be accessed online at: https://www.protocols.io/private/8F324FC92A0AllEDBAlE0A58A9FEAC02

### Equipment

□ 200 μL and 1000 μL pipets
□ Vacuum chamber & Vacuum source reaching at least 0.8 bar of vacuum
□ Oven reaching at least 80°C
□ Magnetic stirring hotplate and magnetic rod
□ 500 mL Beaker
□ 50-100 mL screw-cap bottles
□ Fume hood
□ Centrifuge for plates reaching
□ Tweezers
□ Thermocycler

### Consumables

□ 12-well plates
□ VWR cover film
□ P200 and P1000 tips
□ Alu foil
□ Petri dish
□ Eppendrof 1.5 mL tubes
□ 0.2 mL PCR tubes

### Reagents

□ PDMS
□ DNase free water
□ Bi-distilled water
□ PVA
□ Ethanol 70%
□ Taq 2X Master Mix
□ Primers

### Mold Fabrication

1. Drill a 5mm diameter hole in the bottom of each well of a 12-well plate
2. Stick the Master Molds to the lid of the 12-well plates using double sided tape and close the plate
3. Prepare the PDMS solution with a 1:10 weight ratio of hardener to base and mix it well. For a 12-well plate, 5 g of PDMS per mold are required
4. Cast the PDMS using a syringe or a cut-off pipet tip through the previously drilled holes. We recommend casting about 2/3 of the wells’ height
5. Place the 12-well plate in a vacuum chamber until outgassing is complete (about 30 minutes)
6. Cure the PDMS in an oven at 80°C for 30-60 minutes
7. Take the molds out of the wells and clean them (steps 19-21)

### PVA Solution

8. Prepare a solution of DNase free water and PVA with a 7:1 mass ratio. Note that a batch of 24 microneedles requires typically 28 g of water for 4 g of PVA

? Pour the desired amount of water in a beaker with a magnetic stirrer, add the corresponding weight of PVA and cover the beaker with an aluminum foil
? Place on a magnetic stirring hotplate at 60°C and 300 rpm until the solution is homogenized and clear
9. Transfer to a screw-cap bottle
10. Before use, let the solution settle until the foam from the stirring has disappeared (around 1 hour but typically overnight)

### Microneelde fabrication

11. Preheat the centrifuge to 40°C
12. Place the PDMS molds in the 12-well plates and tape the bottom holes with a VWR cover film
13. Pipet 750 μL of the PVA solution in each mold then close the well plates
14. Centrifuge the plates at 4000 rpm and 40°C for 20 minutes
15. Pipet enough PVA in each molds (maximum 750 μL) to fill the imprints. Note that overfilling the molds will lead to much longer drying times
16. Place the plates without a lid under a fume hood and let dry for 36 hours
17. Use tweezers to extract the Microneedle patches from the molds
18. Remove the molds from the well plates to clean them

### Mold Cleaning

19. Place the PDMS molds with a magnetic stirrer in a beaker, pour bi-distilled water until it covers the molds and cover the beaker with an aluminum foil
20. Place the beaker on a magnetic stirrer hotplate for 1 hour at 200-250 rpm and 80-90°C
21. Take the molds out, rinse them generously with ethanol, shake off the excess and place them under a fume hood until they are dry (about 15 min)

### DNA Extraction

22. Puncture the leaf sample with a Microneedle patch

? Select and stack 2-3 fresh leaves (young leaves are preferred)
? Place a Microneedle patch on the leaf in an area with the least veins possible
? Press forcefully and evenly on the whole patch to puncture the leaf and maintain pressure for 10-15 seconds
? Remove the Microneedle patch and place it on a clean surface (eg: Petri dish)
23. Elute the DNA by pipetting DNase free water on the Microneedle patch:

? For increased DNA concentration, elute with 50 μL
? For increased DNA yield, elute with 100 μL
? Pipet the DNase free water on the Microneedles taking care to wet the whole surface
? Pipet up and down until bubbles form in the water. Avoid leaving the water for too long or the Microneedle patch will dissolve, complicating the recovery of the eluted DNA
? Recover the eluate in an Eppendorf tube

### DNA Amplification

24. Prepare a 50 μL PCR reaction mix:

25. Load the sample in a thermocycler and run the following program:

| Reagent | Volume |  |
| --- | --- | --- |
|  | Microneedle extractions | Qiagen extractions |
| Taq 2X Master Mix | 25 µL | 25 µL |
| Forward Primer | 1 µL | 1 µL |
| Reverse Primer | 1 µL | 1 µL |
| Water | 16 µL | 21 µL |
| DNA Template | 7 µL | 2 µL |

25. Load the sample in a thermocycler and run the following program:
| Cycles | Step | Temperature | Time |
| --- | --- | --- | --- |
| 1x | Initial Denaturation | 95°C | 2 m |
|  | Denaturation | 95°C | 30 s |
| 45x | Annealing | 54°C | 40 s |
|  | Extension | 68°C | 40 s |
| 1x | Final Extension | 68°C | 10 m |
| 1x | Hold | 4°C | ∞ |

## Appendix 3. PCR Primers

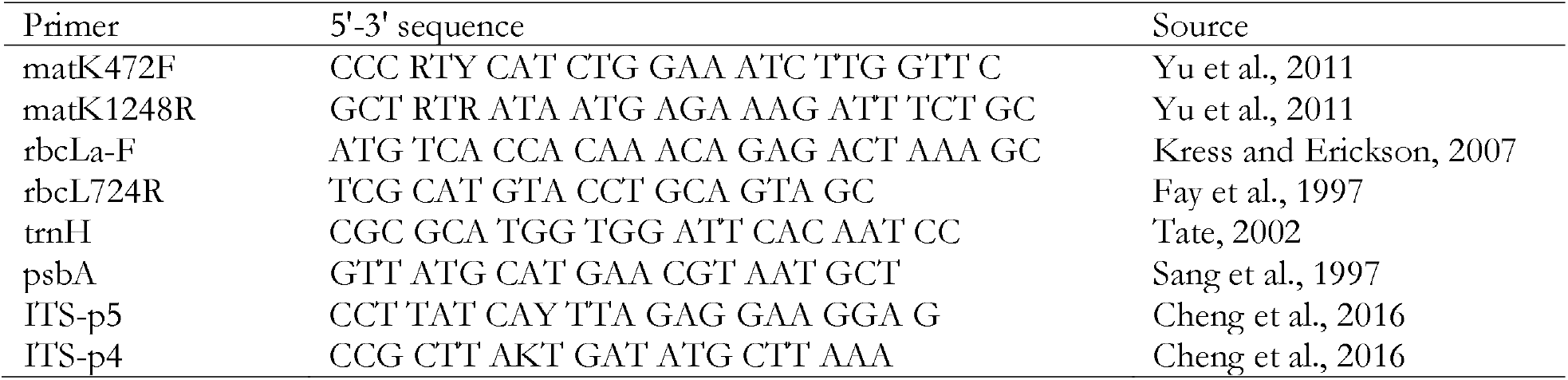

## Notes

### Competing Interest Statement

The authors have declared no competing interest.

